# Percentage of Foveal versus Total Macular Geographic Atrophy as a Predictor of Visual Acuity in Age-related Macular Degeneration

**DOI:** 10.1101/544254

**Authors:** Saghar Bagheri, Ines Lains, Rebecca Silverman, Ivana Kim, Dean Eliott, Rufino Silva, John Miller, Deeba Husain, Joan W Miller, Leonide Saad, Demetrios Vavvas

## Abstract

**Objectives:** To investigate the relationship between visual acuity (VA), total area of geographic atrophy (GA) and percentage of foveal GA.

**Methods:** Multicenter, retrospective cross-sectional study of patients with GA due to age-related macular degeneration. Demographics, VA, fundus autofluorescence (FAF) and optical coherence tomography (OCT) images were collected. Using FAF images aided by OCT, foveal sparing status, GA pattern, total GA size, and percentage of GA covering the foveal area - area within a 1.5 mm diameter circle centered on the fovea centralis - were assessed. Univariable and multiple linear regression analyses were performed.

**Results:** 54 eyes (mean age 78.7 ±7.7 (SD), 60.0% female) were studied. Mean VA was 0.8 ± 0.6 logMAR, mean total GA 8.8 ± 6.7 mm^2^ and mean percentage of foveal GA was 71.5 ± 30.9%. Of all assessed eyes, 48.2% (n = 26) presented with multifocal GA, and 18.5% (n = 10) had foveal sparing. Multiple regression analysis revealed that, controlling for age and GA pattern, the percentage of foveal GA presented a statistically significant association with VA (ß = 0.41, P = 0.004). No significant associations were observed with mean total GA size, while controlling for the same variables (ß = 0.010, P = 0.440).

**Conclusion:** Percentage of foveal GA was significantly associated with VA impairment, while the same was not verified for total GA area. These findings suggest that percentage of foveal GA may represent a more useful tool for assessing the impact of GA on VA. Further validation is needed in larger cohorts.

## INTRODUCTION

Age-related macular degeneration (AMD) is the leading cause of visual disability in elderly patients in industrialized countries.^1^ Geographic Atrophy (GA) represents the late stage of dry AMD, and it is characterized by the irreversible loss of macular retinal tissue, retinal pigment epithelium (RPE), and choriocapillaris.^2^ Although this process occurs in a slowly progressive way, it causes decreases in central vision over time^3^, which rapidly accelerates when GA covers the foveal center. GA is responsible for severe vision loss in approximately 20% of all patients with AMD, may affect up to 22% of the population in 90-year-old people ^2–4^, and more than 8 million people are affected worldwide^2, 4^. For not well understood reasons, atrophic macular diseases such as GA due to dry AMD can spare the foveal center until late in the disease course and the so called foveal sparing has been reported in about 20% of representative GA populations enrolled in clinical trials ^5^.

Color fundus photography, fundus autofluorescence (FAF) and optical coherence tomography (OCT) imaging can be used to identify and follow GA lesions. However, FAF is considered by most to be the imaging of choice that allows for a sharp discrimination of a lesion’s boundaries. This is primarily because FAF provides a good visualization of the high contrast between atrophic (hypofluorescent) and normal areas ^4, 6^. On OCT, GA is typically characterized by the presence of thinning of the hyperreflective external bands due to attenuation/loss of the photoreceptors, ellipsoid zone and retinal pigment epithelium (RPE)/Bruch’s complex; as well as the presence of deeper hyper-reflectivity in the sub-RPE layers due to increased laser light penetration through the atrophic RPE ^2^. The total area of GA lesions is often used as an indicator of severity in late-stage dry AMD. However, this measure does not readily predict residual visual acuity (VA) nor VA decline rates ^7^. Foveal sparing status has been shown to correlate better with VA than total GA size, nevertheless, its binary nature prevents it from being used to quantify the continuous shrinking of the spared foveal area and the worsening of VA over time ^8^.

To explore more sensitive anatomical predictors of VA in GA, we defined and analyzed the percentage of foveal GA and its association with VA. This may lead to more accurate outcome measures for clinical trials as well as for patient counseling.

## METHODS

### Study Design

This study is a multicenter, retrospective cross-sectional study. The research protocol was conducted in accordance with Health Insurance Portability and Accountability Act requirements and the tenets of the Declaration of Helsinki. The Institutional Review Boards of MEE and of the Coimbra University Hospital approved this study.

### Study Population and Study Protocol

We identified and retrospectively reviewed the medical records and images of eyes with GA. We adopted the most recent AREDS definitions ^9^, namely that geographic atrophy is present if the lesion has a diameter of 433 μm or more (AREDS circle I-2) and has at least 2 of the following features: absence of retinal pigment epithelium (RPE) pigment, circular shape, or sharp margins.

Subjects from two centers were considered. At MEE, we identified patients seen between September 2011 to June 2017 as part of the AMD biomarkers study and from the attending clinic (DGV). From Portugal, we considered subjects participating in the AMD biomarkers study, developed by the Faculty of Medicine, University of Coimbra, in collaboration with the Association for Innovation and Biomedical Research on Light and Image (AIBILI) and the “Centro Hospitalar e Universitário de Coimbra”, Coimbra, Portugal.

For all considered subjects, exclusion criteria included: GA with CNVM; diagnosis of any other vitreoretinal disease, active uveitis or ocular infection; significant media opacities that precluded the observation of the ocular fundus; refractive error equal to or greater than 6 diopters of spherical equivalent; history of any ocular surgery or intra-ocular procedure such as laser or intravitreal injections within 90 days prior to enrollment; and diagnosis of diabetes mellitus. Additionally, only eyes with both FAF and OCT images according to a predefined protocol, available on at least one visit were considered for this study. For FAF, we considered eyes with high resolution 30° FAF, centered on the fovea. For OCT, we used high resolution 30° spectral domain optical coherence tomography (SD-OCT).

For the final included eyes, we reviewed medical records and collected the following information: age, gender, smoking status, AREDS supplementation and Snellen VA at the same date of the considered images.

### Imaging analysis

We reviewed FAF and OCT images of the eyes considered for this study. The fovea was defined by a 1.5 mm diameter circle area of 1.77 mm^2^, centered on the fovea centralis (Figure 1). Using the Heidelberg built-in calipers, two masked graders (S.B., R.S.) independently measured the total GA area and the percentage of the foveal area covered by GA on the FAF images. The same graders also assessed foveal sparing status. For this grading process, OCT images were used in parallel to help determine the location of the umbo/fovea centralis and the GA areas in a multimodal approach^10^. For analysis, average values of the two graders were used, except when values disagreed by more than 10%, in which case a third grader (I.L.) was used for adjudication. Furthermore, we graded for foveal sparing status and GA pattern (focal or multifocal) in addition to collecting demographic information on age, gender, study eye, smoking status and AREDS supplementation.

**Figure 1.**
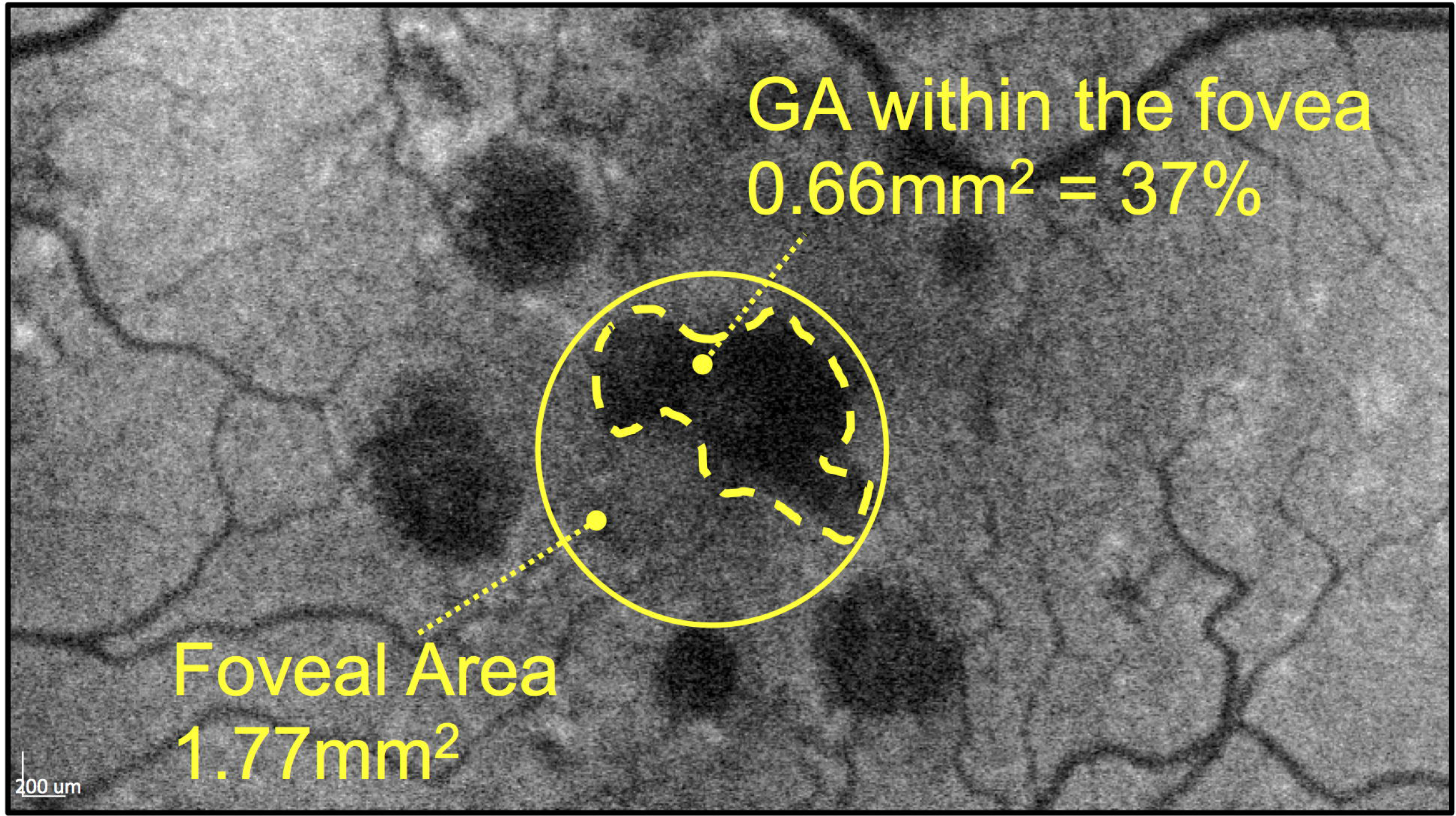
Percentage of foveal geographic atrophy (GA) Representative blue autofluorescence fundus image outlining geographic atrophy (GA) within the foveal area, defined by a 1.5 mm diameter circle centered on the fovea centralis which has been determined with the use of optical coherence tomography (OCT) images.

### Statistical and Data Analysis

Traditional descriptive methods such as mean and standard deviation for continuous variables, and percentages for dichotomous/categorical variables were used to describe the clinical and demographic characteristics of the study population.

Regarding the inclusion of two eyes of the same patient for some cases, our statistical assessments were performed using multilevel mixed effect models. By definition, these models are appropriate for research designs in which data for participants are organized at more than 1 level (ie, nested data). In this study, the units of analysis were considered the eyes (at a lower level), which are nested within patients that represent the contextual/aggregate units (at a higher level)^11^.

Univariate analyses were initially performed for all the potential confounders such as age and GA pattern, and all variables with a P value ≤ 0.250 were included in the initial multiple model. A backwards elimination procedure was then performed to achieve the multivariable models presented. For both univariate and multivariate analyses, we report *P* values and beta coefficients. The beta coefficients represent the change in the outcome variable for 1 unit of change in the predictor variable (while holding other predictors in the model constant, in the case of multivariate analyses) ^12^. This means, for example, given a continuous variable such as age, beta coefficients represent the change in visual acuity per year increase in age. For binomial variables, such as smoking, AREDS supplementation or foveal sparing, their absence was considered the reference term, so beta coefficients refer to the change in their presence. The reference terms for study eye was the right eye, for GA pattern unifocal GA and for gender was the female gender.

All statistics were performed using Stata version 14.1 (StataCorp LP, College Station, Texas, USA) and P values < 0.05 were considered statistically significant.

## RESULTS

### Study Population

We included 54 eyes from 35 patients (mean age 78.7 ± 7.7 years, 60.0% female (n = 21)) with GA due to non-neovascular AMD. Mean VA was 0.81 (20/129 Snellen eq.) ± 0.63 [range, 0 to 2.60] logMAR, mean total GA 8.79 ± 6.66 [0.84-25.36] mm^2^, mean percentage of foveal GA was 71.53 ± 30.94 [0-100] %. 48.15 % (n = 26) of assessed eyes presented with multifocal GA, and 18.52% (n = 10) had foveal sparing (see demographics in Table 1).

**Table 1.** Demographic and Clinical Characteristics of the Included Study Eyes

|  |  |
| --- | --- |
| Age at date of imaging in years (n=54) |  |
| Mean ± SD (range) | 78.7 ± 7.7 (62 to 96) |
| Gender (n=35) |  |
| Female | 21 (60.0%) |
| Male | 14 (40.0%) |
| Smoking (n=30) |  |
| no | 27 (90.0%) |
| yes | 3 (10.0%) |
| AREDS (n=38) |  |
| no | 14 (36.8%) |
| yes | 24 (63.2%) |
| Study Eye (n=54) |  |
| OD | 29 (53.7%) |
| OS | 25 (46.3%) |
| GA pattern (n=54) |  |
| unifocal | 28 (51.9%) |
| multifocal | 26 (48.2%) |
| Foveal Sparing (n=54) |  |
| no | 44 (81.5%) |
| yes | 10 (18.5%) |
| VA in logmar (n=54) |  |
| Mean ± SD (range) | 0.8 ± 0.6 (0 to 2.6) |

GA = Geographic Atrophy; VA = Visual Acuity.

In all eyes, SD-OCT images allowed a clear identification of the umbo/fovea centralis as well as the GA lesion borders. Figure 1 presents an example of measurement of percentage of foveal GA. The results of the univariate analysis considering all variables of interest and their association with VA is shown in Table 2.

**Table 2.** Univariable Linear Regression Analysis Considering VA as the Outcome

| | $\beta$ | *P* value | 95% CI |
| --- | --- | --- | --- |
| Age, years | 0.017 | 0.113 | [-0.00409, 0.038529] |
| Gender<sup>a</sup> | -0.285 | 0.097 | [-0.6215504, 0.0512273] |
| Study Eye<sup>b</sup> | -0.226 | 0.178 | [-0.5543471, -0.102713] |
| Smoking<sup>c</sup> | -0.319 | 0.398 | [-1.058263, 0.4205226] |
| AREDS<sup>c</sup> | -0.412 | 0.051 | [-0.8265572, -0.0018414] |
| GA pattern<sup>d</sup> | -0.507 | 0.001* | [-0.8118193, -0.2025564] |
| Foveal sparing<sup>c</sup> | -0.539 | 0.009* | [-0.9440415, -0.134384] |
| Total GA (mm<sup>2</sup>) | 0.024 | 0.054 | [-0.0004198, 0.048712] |
| % of foveal GA | 0.536 | 0.000* | [0.2625702, 0.8104187] |

VA = Visual Acuity; CI = confidence interval; GA = Geographic Atrophy.
*P* values < 0.05 are noted by an asterisk.
<sup>a</sup> Female gender considered the reference term.
<sup>b</sup> Right eye considered the reference term.
<sup>c</sup> Reference term considered as the absence of these variables.
<sup>d</sup> Unifocal Geographic Atrophy considered the reference term.

### Percentage of foveal GA

The mean percentage of foveal GA was statistically significantly associated with VA in univariate analysis (ß = 0.54, P < 0.001) (Table 2). This association remained significant on multivariate analysis, controlling for age and GA pattern (ß = 0.41, P = 0.004).

### Total macular GA

Univariate or multivariate analysis for total macular GA revealed that there were no statistically significant associations with VA (ß = 0.02, P = 0.054, for univariate), (ß = 0.010, P = 0.440 for multivariate).

### GA pattern & Foveal Sparing

GA pattern presented a statistically significant association with VA (ß = −0.5071879, P = 0.001) and so did foveal sparing (ß = −0.5392127, P = 0.009).

## DISCUSSION

We present a retrospective, cross-sectional study of 54 eyes diagnosed with GA due to non-neovascular AMD, in which we used FAF and SD-OCT to examine the associations of percentage of foveal GA and total macular GA lesion size with VA. Our results revealed that, after accounting for potential confounders such as age and GA pattern, the percentage of foveal GA was significantly associated with VA, while the same was not observed for total GA lesion size.

In GA clinical studies, the most common outcome measures for GA are changes in total GA, changes in square root GA, or other phenotypic refinements ^13, 14^. As our results show, total GA poorly correlates with VA, and potentially patients’ overall quality of life. This finding is in agreement with previously published literature, which showed no relationship of total GA size with VA and has been investigated by multiple groups ^8, 15^.

Efforts have been made to study the association between VA and the distance between the edges of GA and the fovea ^15, 16^, or to examine residual visual function in the presence of foveal sparing lesions ^15–17^. Foveal sparing status has been shown to have a stronger correlation with VA than total GA size, however, it does not quantify the extent to which the foveal area is affected nor the worsening of VA over time since it only measures presence or absence of geographic atrophy in the anatomic foveola centralis ^8, 15, 16^. A recent investigation of the associations of VA with total GA size as well as foveal sparing status in 65 eyes found no relationship between VA and total GA size as well as foveal island size ^18^. The same group also evaluated the width of the bridge - defined as the minimal linear dimension of intact RPE located within the residual foveal island - and only found a suggestion of a positive relationship in the range of 300 to 550 μm of bridge width and no relationship at all outside of this range leading to the conclusion that this measurement might not be an ideal outcome parameter for GA clinical trials.

Our study results suggest that using the percentage of foveal GA is potentially a more sensitive outcome parameter for association with visual acuity. Our study however, is limited by its modest size and retrospective design. As such, our results should be validated in larger, more representative populations, before changes in percentage of foveal GA can be used more widely in clinical trials or clinical practice. Further studies should examine more precise evaluation of affected areas as well as evaluate the progression rate of percentage of foveal GA over time and examine predictive ability of such tool on future VA changes.

In conclusion, here we propose for the first time the use of percentage of foveal GA as a possible predictor of VA in GA. Our data suggests that such a measure may have a stronger association with VA impairment than total GA size. Therefore, with future research, it might represent a better tool to measure VA decline over time compared to the foveal sparing status.

## Acknowledgments

None.

## Ethical Approval

The research protocol was conducted in accordance with Health Insurance Portability and Accountability Act requirements and the tenets of the Declaration of Helsinki. The Institutional Review Boards of MEE and of the Coimbra University Hospital approved this study.

